# Improved selection of canonical proteins for reference proteomes

**DOI:** 10.1101/2024.03.04.583387

**Authors:** Giuseppe Insana, Maria J. Martin, William R. Pearson

**Affiliations:** European Molecular Biology Laboratory, European Bioinformatics Institute (EMBL-EBI), Wellcome Genome Campus, Hinxton, CB10 1SD; Dept. of Biochemistry and Molecular Genetics, U. of Virginia School of Medicine, Charlottesville, VA 22908, USA

## Abstract

The “canonical” protein sets distributed by UniProt are widely used for similarity searching, and functional and structural annotation. For many investigators, canonical sequences are the only version of a protein examined. However, higher eukaryotes often encode multiple isoforms of a protein from a single gene. For unreviewed (UniProtKB/TrEMBL) protein sequences, the longest sequence in a Gene-Centric group is chosen as canonical. This choice can create inconsistencies, selecting *>*95% identical orthologs with dramatically different lengths, which is biologically unlikely. We describe the ortho2tree pipeline, which examines Reference Proteome canonical and isoform sequences from sets of orthologous proteins, builds multiple alignments, constructs gap-distance trees, and identifies low-cost clades of isoforms with similar lengths. After examining 140,000 proteins from eight mammals in UniProtKB release 2022 05, ortho2tree proposed 7,804 canonical changes for release 2023 01, while confirming 53,434 canonicals. Gap distributions for isoforms selected by ortho2tree are similar to those in bacterial and yeast alignments, organisms unaffected by isoform selection, suggesting ortho2tree canonicals more accurately reflect genuine biological variation. 82% of ortho2tree proposed-changes agreed with MANE; for confirmed canonicals, 92% agreed with MANE. Ortho2tree can improve canonical assignment among orthologous sequences that are more than 60% identical, a group that includes vertebrates and higher plants.

## Introduction

Comprehensive, well-curated protein sequence databases have transformed molecular biology. Since the early discoveries of homology between viral oncogenes and protein kinases [1] and also growth factor receptors [2], protein sequence similarity has become the most common and most effective strategy for inferring protein function. Indeed, for non-model organisms, 80–100% of functional annotations in the Gene Ontology resource [3] are inferred solely from sequence data. Because it is so effective, the BLAST [4, 5] similarity searching program is one of the most highly cited methods in the biomedical literature.

Similarity searches are usually done against reference protein databases. The UniProt Reference Proteomes (release 2023 05, November 2023) provide 24,004 protein sets, about half from viruses (11,951), another 9,793 from prokaryotes (bacteria and archaea), and 2,260 from Eukaryotes [6]. For this latter set, about a third are from vertebrates (623) and higher plants (243), groups of organisms that can produce multiple protein isoforms from a single gene. When multiple isoforms are produced by a single gene, the Reference Proteome set selects, using a Gene-Centric approach, a single isoform as “canonical”, i.e., the representative sequence for all the isoforms transcribed from that gene. The selected canonical proteins are distributed for sequence similarity searching, are functionally annotated, and are characterized for domain content [7]. The focus on canonical sequences for functional and structural annotation makes canonical isoform selection critical.

UniProtKB/Swiss-Prot sequences in the UniProt References proteomes are manually curated [6], but the vast majority of non-model organism sequences are annotated automatically, and, until UniProt release 2023 01, the longest isoform was chosen as the canonical sequence. This longest-canonical rule can produce some very non-biological results (Fig. 1). For example, in comparisons of mouse and rat proteins, there are 25 best pairwise alignments that are more than 99% identical (4 that are 100% identical) but have gaps of more than 50 residues. Additionally, in release 2022 05 of the UniProt reference proteomes, the best alignment of mouse bis-phosphate mutase (UniProt accession P15327, 259 amino-acids) to canonical rat proteins is a 96% identical alignment with Q7TP58, a 395 amino-acid protein, with a 104 insertion in the rat protein. However, a rat isoform (Q6P6G4, 258 amino-acids) matches the mouse enzyme without any gaps.

**Figure 1:**
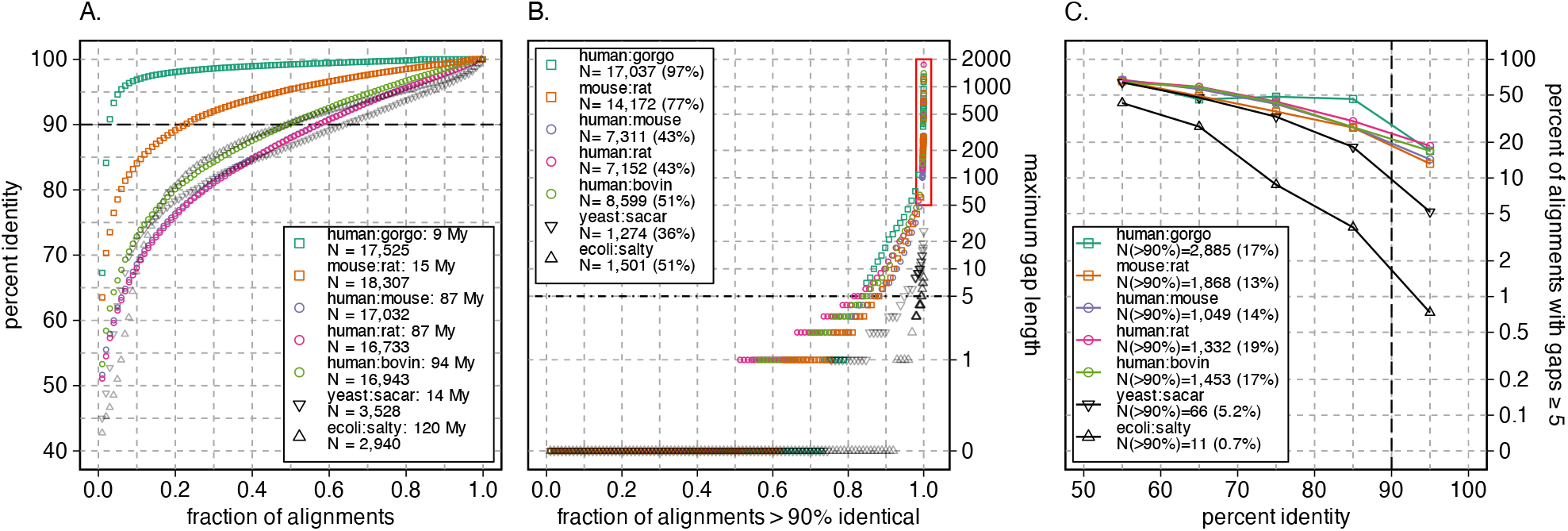
Identity and gap lengths for pairs of mammalian, yeast, and bacterial proteomes. The complete canonical reference proteome of human (UniProt release 2022 05, 20,594 sequences) was compared to the canonical reference proteome of gorilla (21,783), mouse (21,968), rat (22,816), cow (23,844), and the canonical mouse proteome was compared to rat. In addition, the canonical *E. coli* proteome (4,402) was compared to *S. typhimurium* (salty) (4,533) and the canonical *S. cerevisiae* (yeast) proteome (6,060) was compared to *S. arboricola* (sacar) (3,653). Searches used the Smith-Waterman algorithm and the VT40 scoring matrix, as described in Materials and Methods. (A) The highest scoring statistically significant hits (E()*<* 10^*−*6^) from each query longer than 100 aa are ranked by percent identity; 100 samples from that ranking are plotted for each pair of proteomes. The legend shows the search pair (e.g. human:gorgo, human:gorilla), the evolutionary distance between the two proteomes, and the number of queries that produced a statistically significant hit. Alignment pairs above the dashed line at 90% identity were used in panel (B). (B) The alignments with *>*90% identity are ranked from shortest to longest maximum gap length and 100 samples, plus all alignments with a maximum gap length of *>*50, and the longest 25 gaps, are plotted. The panel (B) legend shows the total number (and percent) of alignments from panel (A) with *>*90% identity. The red rectangle highlights alignments with gaps *≥*50 aa. (C) Percent of alignments with a gap *≥*5 residues versus percent identity. The legend shows the number of alignments with a gap *≥*5 residues for alignments that are *>*90% identical; this number corresponds to the points plotted at 95% identity in panel (C).

Even a small fraction of incorrect canonical sequences can dramatically reduce similarity search sensitivity, in part by producing artifactual domain architectures [8]. These artifactual architectures can provide high-scoring alignment regions in nonhomologous proteins that contaminate models produced by iterative searches [5, 9, 10],

The problem of length differences in closely related protein sequences has been recognized for some time [11, 12, 13], and strategies have been developed to select representative isoforms to be canonical. The PALO strategy [11] builds a set of protein isoforms that are most similar in length for evolutionary rate studies. This has the advantage that it is computationally efficient, but it can group together isoforms that have different sets of exons that add up to similar lengths.

The APPRIS database [12, 14] annotates isoforms using a combination of structural, domain, functional sites, and conservation information. In addition to annotating a principal isoform, the simple identification of a “canonical” isoform, APPRIS also annotates additional isoforms as likely to be biologically significant. APPRIS currently annotates 9 vertebrate genome assemblies, as well as *D. melanogaster* and *C. elegans* (appris.bioinfo.cnio.es/). In addition to APPRIS, MANE [13], a joint NCBI and EMBL-EBI annotation group, has produced a reference set of human transcripts and corresponding protein isoforms for human genes. The MANE-select protein set is annotated using a set of analysis pipelines and manual curation, but is only available for human proteins.

In this paper, we describe the ortho2tree protein analysis pipeline, which uses a gap-distance tree-based strategy for identifying preferred canonical isoforms, and show that it identifies canonical protein isoforms with gap distributions similar to those seen in bacteria and yeast alignments. Bacteria do not produce protein isoforms, and fewer than 1% of yeast proteins have isoforms, so these protein alignments serve as a reference for the types of gaps produced by evolutionary divergence. Thus, the alignment similarity between ortho2tree selected isoforms and bacterial/yeast protein alignments suggests that the selected isoforms reflect genuine biological, rather than artifactual isoform-selection, variation.

Like the PALO [11] strategy that seeks to reduce length variation, ortho2tree begins with the assumption that orthologous closely related proteins, e.g. proteins that have diverged from a common ancestor in the past 100–200 million years (My), should align with very few long gaps. Ortho2tree begins with sets of orthologous proteins and their isoforms, produces a multiple sequence alignment from those proteins, and then creates a phenogram based only on the gap-based distances between the orthologs. This tree is then examined to find low-cost (similar length) clades containing sequences from the distinct proteomes. Beginning with 8 mammals in UniProt release 2023 01, the ortho2tree approach has been used to modify canonical isoform for the UniProt reference proteome set. In release 2024 01, it modified canonical isoform assignments for 49,252 proteins from 35 mammals. With new strategies for comprehensive ortholog identification, we believe that the ortho2tree pipeline can be used to improve canonical isoform assignments for proteomes with orthologous sequences that share more than 50% sequence identity. These groups include vertebrate and plant proteomes. We have begun updating plant canonical sequence assignments with Uniprot Release 2024 02, and we look forward to extending the pipeline to vertebrate proteomes.

## Materials and Methods

### Similarity searches

Sequence databases were obtained from the UniProt Reference Proteomes site (ftp.ebi.ac.uk/pub/databases/uniprot/previous_releases/release-2022_05).

Sequence databases were downloaded from the UniProt Knowledgebase in FASTA format, and then low-complexity regions were soft-masked using the pseg [15] program. Similarity searches were done using programs from the FASTA package [16] (version 36.3.8i, May, 2023), using the VT40 scoring matrix [17], which targets alignments that are more than 60% identical [18]. fasta36 and ssearch36 searches used the -M8CBl output option, which provides a compact summary of each alignment, including the BTOP alignment encoding used by the BLAST programs [19]. Search results with the alignment encodings were parsed to summarize the gaps and gap length statistics. Divergence time estimates were obtained from timetree.org [20, 21].

#### The ortho2tree pipeline

The ortho2tree analysis pipeline goes through five steps, outlined in Fig. 2, to gather orthologous isoforms, align them using muscle [23], build a gap-based evolutionary tree, find low-cost diverse clades, and rank the low-cost clades.

**Figure 2:**
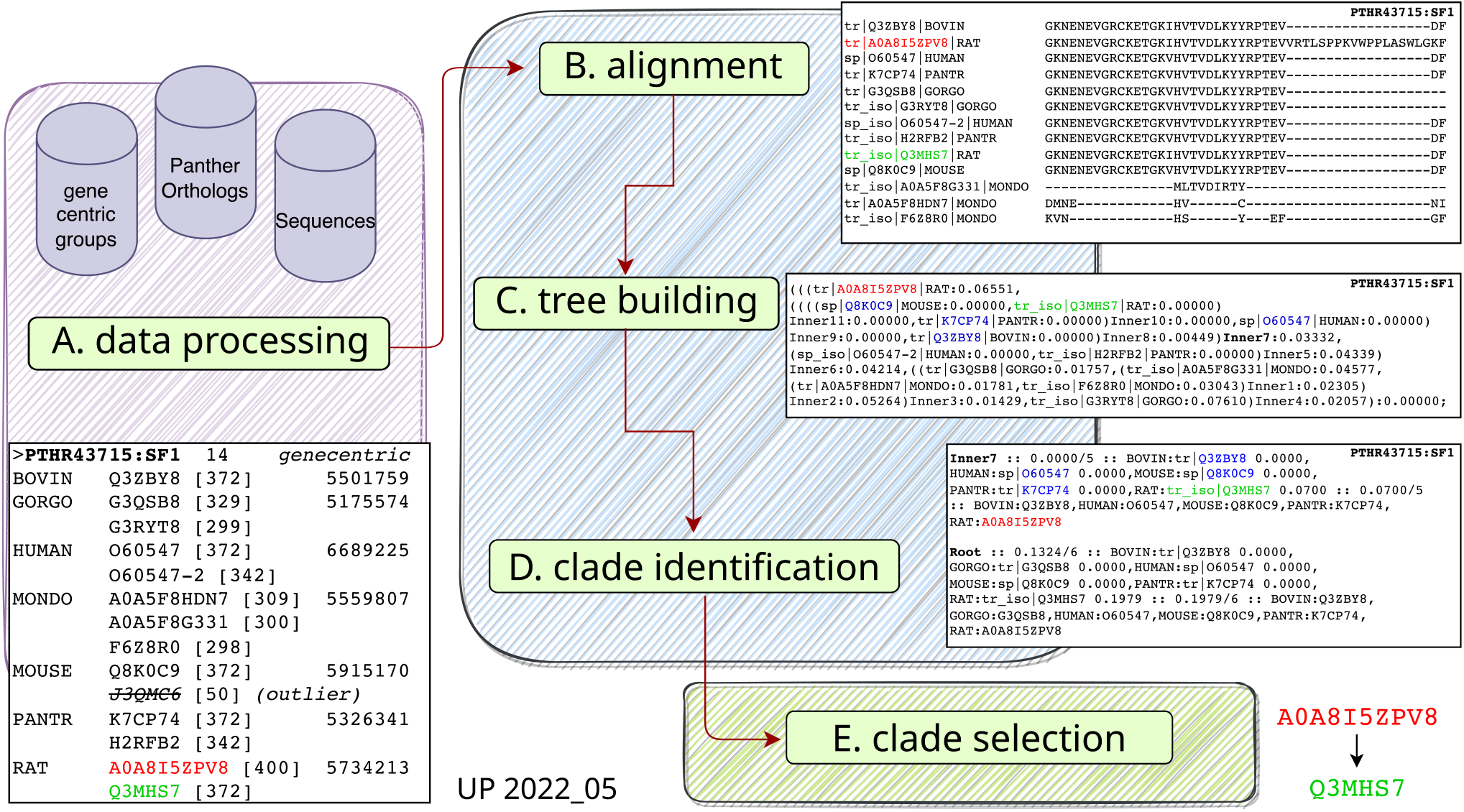
The ortho2tree analysis pipeline. (A) Gene-Centric groups for Panther orthologs [22] are used to extract orthologous canonical and isoform sequences. Panther Ortholog IDs are used to group genes among organisms; Gene-Centric groups are used to identify canonical and isoform sequences associated with each Panther Ortholog set. (B) Orthologous canonical proteins and orthologs are aligned using muscle [23]. (C) A gap-distance tree (see Methods) is constructed using the BioPython3.8 [24] functions DistanceCalculator, DistanceMatrix, and DistanceTreeConstructor. (D) A heuristic clade search script examines the gap-distance tree and identifies low-cost clades with sequences from different proteomes. (E) The list of low-cost clades is scored based on the number of proteomes, the cost of the clade, the number of UniProtKB/Swiss-Prot entries from human or mouse, and length consistency. The highest scoring clades from each distinct gene set is then plotted on a tree (Fig. 3).

(A) Ortho2tree begins by identifying the sequences associated with each Panther orthogroup family in each of the target proteomes. Panther assigns the canonical UniProt accession to an orthogroup; this set of canonical accessions is expanded to a complete set of canonical and isoform accessions using the Gene-Centric mapping (Fig. 2). The pipeline can retrieve protein sequences using direct access to the UniProt databases or using the UniProt web API [25]. This process collects all the canonical proteins, and their isoforms, for each of the target proteomes for the 23,878 Panther orthogroups present in the analyzed mammals (Panther v. 17, Table 1). These steps are illustrated in Fig. 2A.

**Table 1:** Yield of Panther orthogroups and UniProt accessions through the Ortho2tree pipeline^*a*^.

|  | groups | total<br>canon. | clade<br>canon. | conf.<br>canon. | prop.<br>change |
| --- | --- | --- | --- | --- | --- |
| Confirmed | 11,010 | 75,628 | 53,300 | 53,300 | – |
| Prop. changes | 5,005 | 35,004 | 26,168 | – | 7,755 |
| Both conf.+prop.<sup>b</sup> | 35 | 338 | 225 | 134 | 49 |
| Skipped<sup>c</sup> | 4,742 | 11,416 | – | – | – |
| No low-cost clades | 3,086 | 19,158 | – | – | – |
| Total | 23,878 | 141,544 | 79,588 | 53,434 | 7,804 |

<sup>a</sup> Analysis based on UP2022\_05.
<sup>b</sup> Groups with multiple low cost clades, both confirming canonicals and proposing a change.
<sup>c</sup> Groups with sequences from < 3 proteomes.

(B) Once a set of sequences is collected for an orthogroup, orthogroups with fewer than three organisms (for the releases shown here) are filtered out (Table 1, “Skipped”). The sequences (canonical and isoform) are then aligned using the muscle [23] multiple sequence alignment program (version 3.8.31, Fig. 2B), using the default scoring parameters, and the -maxiters 2 -diags command line options. Next, a “gap-distance” matrix is calculated from the multiple sequence alignment using a modified version of the BioPython (v3.8, [24]) Bio.Phylo.TreeConstruction DistanceCalculator function that only counts gapped residues. Thus, two sequences that are 90% identical but align end-to-end with no gaps, have a distance of zero; two sequences that with 100% identical amino-acids, but with a 5 residue gap in the alignment, would have a distance of 5/alignment-length.

(C) The pairwise gap-distances are then used to calculate a gap-based Neighbor-Joining trees [26] using the DistanceTreeConstructor function of BioPython (Fig. 2C).

(D) Each Neighbor-Joining tree is then scanned to identify low-cost clades that contain one sequence from as many proteomes as possible (Fig. 2D). Each low-cost clade contains only one protein sequence from each of the proteomes. Examples of these clades can be seen in Fig. 3.

**Figure 3:**
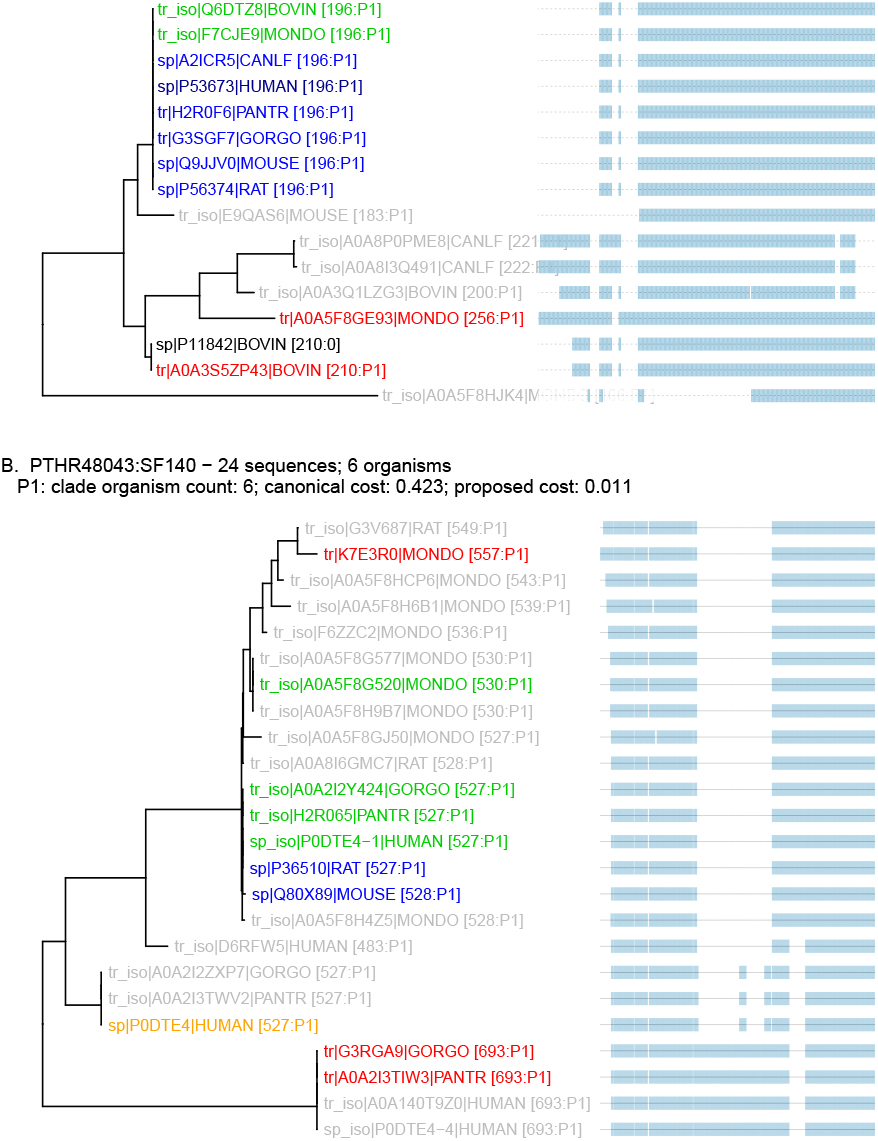
Phenograms highlighting selected clades for two Panther orthogroups: (A) PTHR11818:SF19 (*β*-crystallin A4); (B) PTHR48043:SF140 (UDP-glucuronosyltransferase, E.C. 2.4.1.17); For each panel, a gap-distance based evolutionary tree and multiple sequence alignment are shown. Each panel also shows the total number of canonical and isoform sequences examined and the number of organisms with a sequence in this orthogroup, followed by a list of proposed replacement (P1) clades and their sizes and costs. In the tree, blue sequences are canonical accessions; the darker sp|P53673|HUMAN sequence in panel (A) was selected by the MANE [13] consortium. Accessions shown in red and orange are proposed to be changed; orange canonicals have been selected by MANE. For both red and orange, there is a corresponding green isoform that is proposed to be canonical. **black** accessions indicate canonical isoforms that are not included in a low-cost clade. Gray accessions indicate non-canonical isoforms. Numbers after each protein accession indicate the length of the sequence, and its assigned clade (P1). This figure was drawn using the ggtree package for ‘R’ [27].

(E) Once low-cost multi-proteome clades have been identified, they are ranked based on: (1) the overall gap-cost of the clade; (2) the number of sequences; (3) the number of canonical sequences; (4) the number of UniProtKB/Swiss-Prot annotated sequences from human and mouse; and (5) length consistency, clades that have lengths very different from the median canonical lengths are down-weighted (Fig. 2E). The highest ranked candidate is chosen first, and additional candidate clades are chosen if they are derived from different sets of Gene-Centric genes (Suppl. Fig. 2).

The “yield” of sequences and orthogroups for the analysis of UniProt release 2022 05 (which was used to select the canonical sequences in release 2023 01) is shown in Table 1. Beginning with 23,878 Panther (v. 17) orthogroups and 141,544 canonical sequences, about one-third of these orthogroups (7,828) were skipped either because they did not have proteins from at least three different proteomes (Table 1, “Skipped”), or because the tree-building/clade-finding analysis did not find a low-cost clade with sequences from at least three proteomes (Table 1, “No low-cost clades”). The remaining 16,050 orthogroups produced low-cost clades with sequences from at least three proteomes. For 11,010 orthogroups, the best clade contained only canonical sequences; for 5,040 orthogroups, a clade containing one or more isoforms had a lower cost (higher score) than a clade containing canonical sequences from the same proteomes; and 35 orthogroups produced a clade that confirmed one set of canonicals while proposing changes for another (e.g., Suppl. Fig. 2). Examples of the phenograms and their corresponding low-cost clades are shown in Fig. 3 for two Panther orthogroups.

The ortho2tree clade ranking strategy seeks a balance of low-cost clades (clades with sequences that align without gaps) and increased proteome coverage. Clades must include isoforms from at least three proteomes, but clades need not contain isoforms from every proteome.

During the development of ortho2tree in 2023, the code used to find low-cost clades was improved. Figs. 3, 4, 5, and Suppl. Figs. 1 – 3, present results using the revised current code. Figs. 6 and 7 present the numbers for the code that was used when the releases were made.

**Figure 4:**
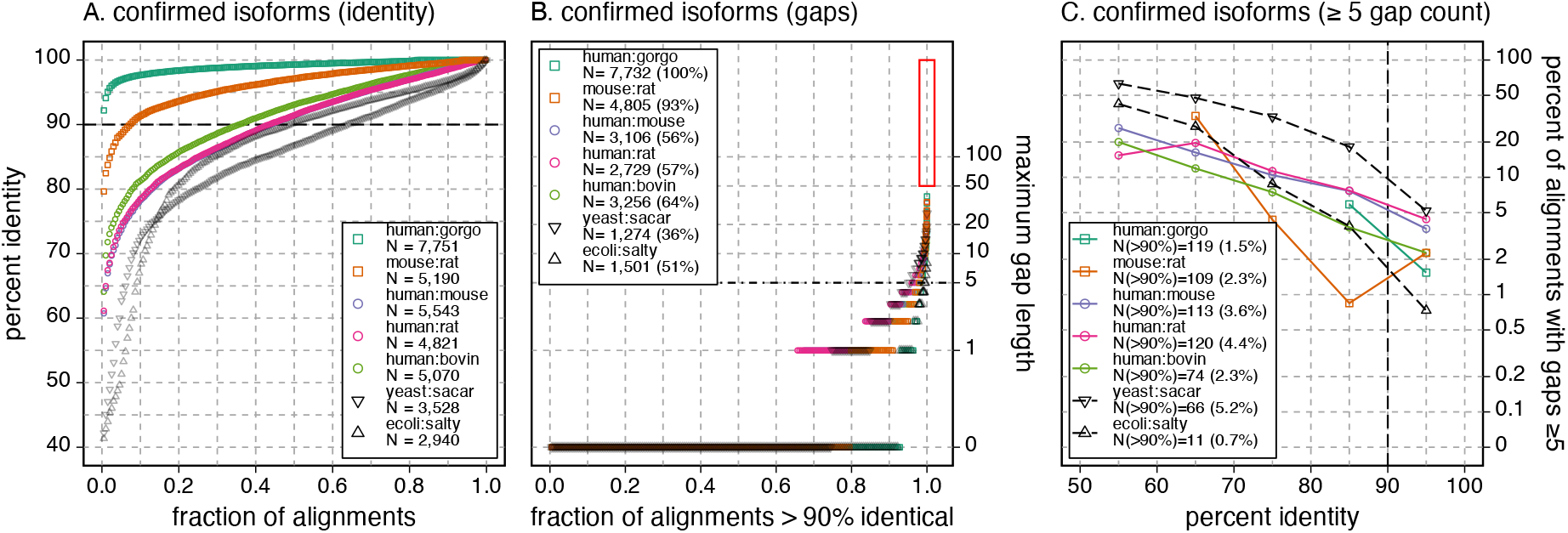
Mammalian confirmed-canonical alignments are similar to bacterial alignments. (A) One hundred samples of percent identity across the 5,244 pairs of sequences for which canonical changes were proposed (circles) and 100 samples from the distribution of identities shown in Fig. 1A for *E. coli* vs *S. typhimurium* and *S. cerevisiae* vs *S. arboricola*. (B) Distribution of gap sizes for the 100 samples shown in (A), plus all alignments with a maximum gap length greater than 50, and the longest 25 gaps. (C) The percentage of alignments with gaps *≥*5 residues versus alignment percent identity for all the pairwise alignments shown in (A). The distributions of ecoli:salty and yeast:sacar alignments in panels A–C replot the data shown in Fig. 1.

**Figure 5:**
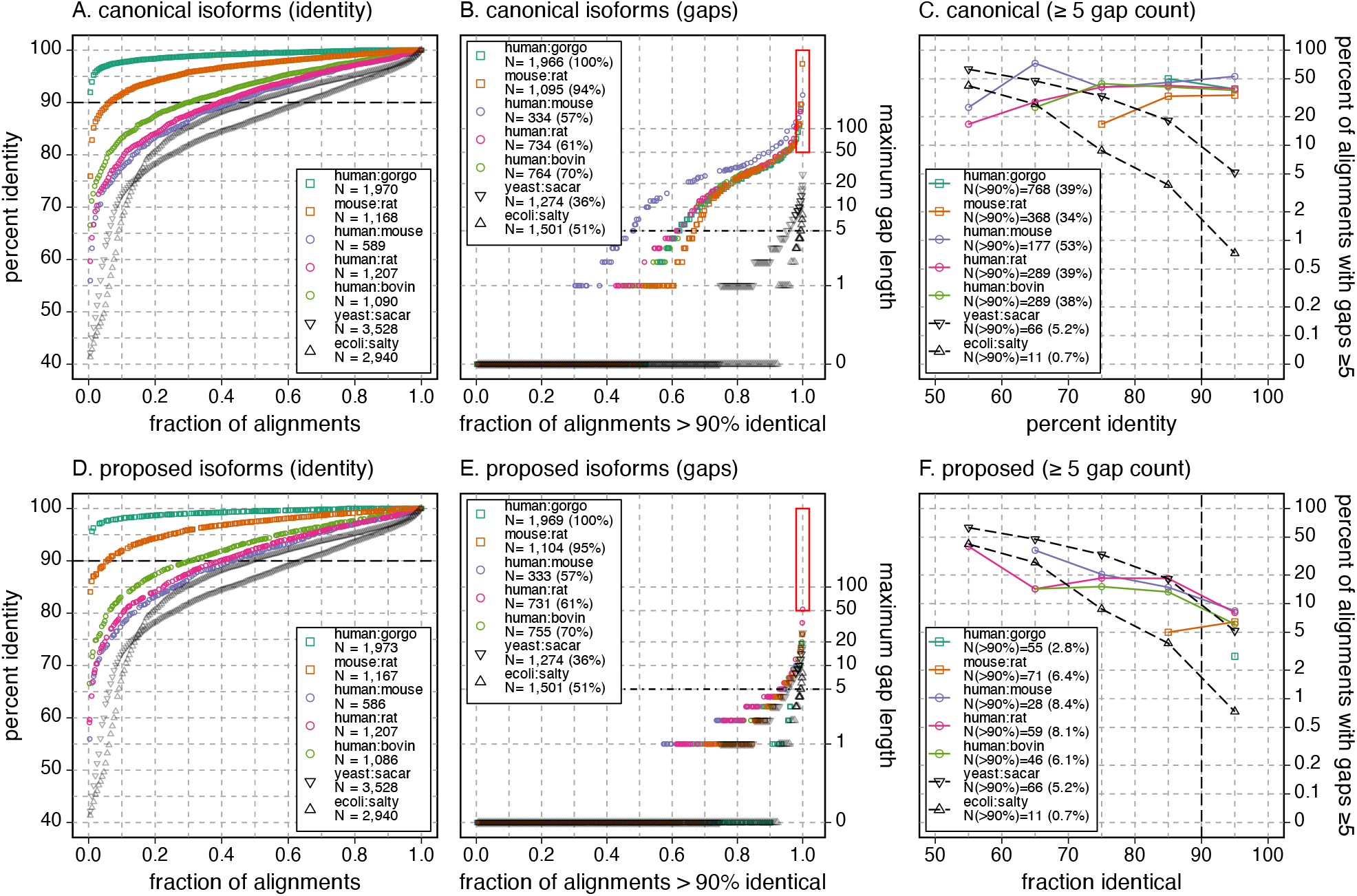
Ortho2tree proposed changes reduce gaps in orthologs. (A) Percent identities for confirmed canonical isoform clades for the proteome pairs shown in Figs. 1 and 4 and the *E. coli* vs *S. typhimurium* and *S. cerevisieae* vs *S. arboricola* comparisons. One hundred samples are shown for each proteome pair. (B) Distribution of the maximum gap sizes for the 100 samples shown in (A), plus alignments with gaps longer than 50 residues and the 25 longest gaps. (C) The percentage of alignments with gaps *≥*5 residues versus alignment percent identity for all the pairwise alignments shown in (A). (D) Percent identity, (E) maximum gap lengths in *>*90% identical alignments, and (F) percent of alignments with gaps *≥*5 residues for the proteins plotted in panels A–C, using the alternative isoforms proposed by ortho2tree. The distributions of ecoli:salty and yeast:sacar in panels A–C and D–F replot the data shown in Fig. 1.

**Figure 6:**
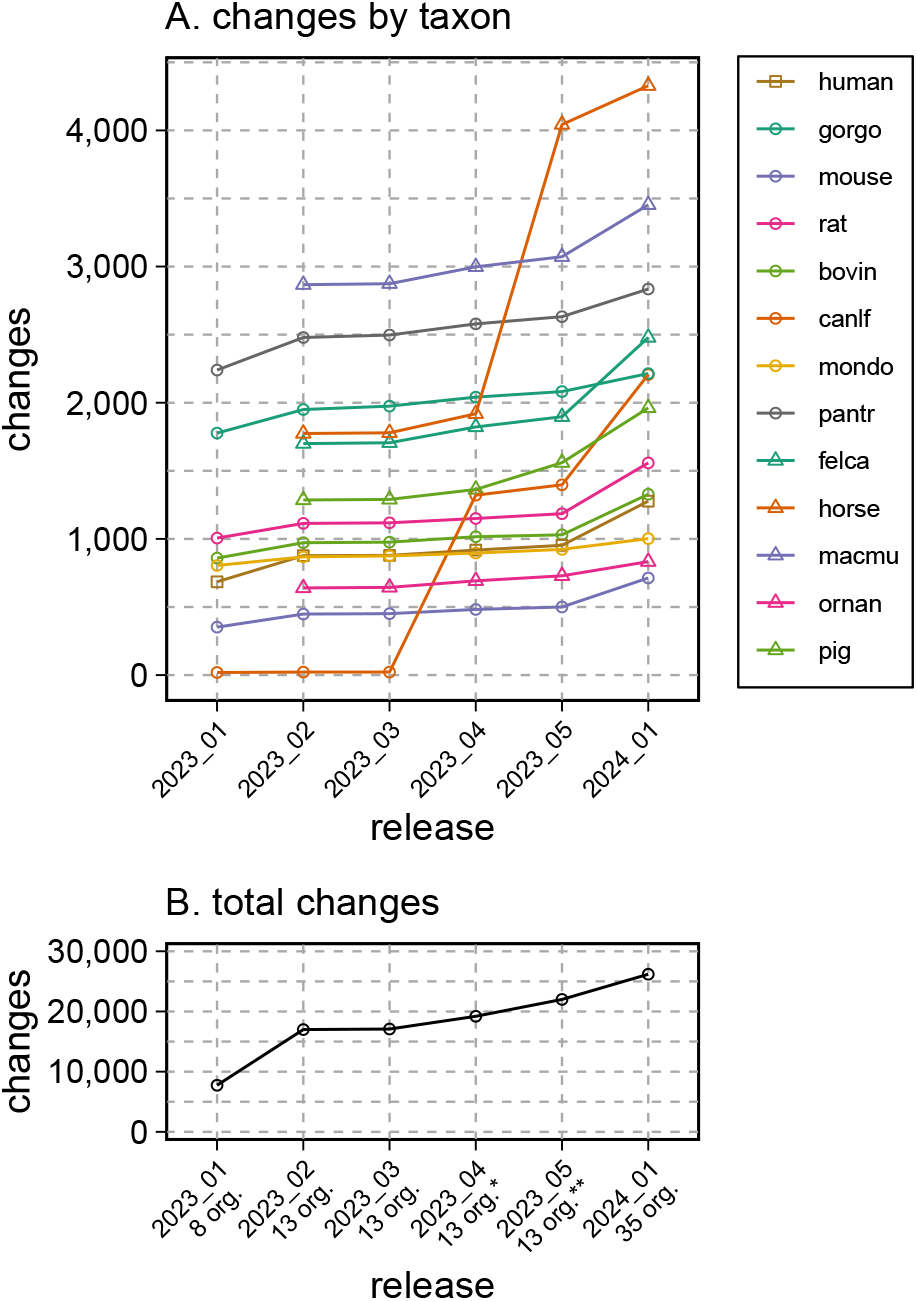
Cumulative numbers of proposed changes for the 13 Panther annotated mammalian proteomes across six UniProt releases. The number of proteomes increases from 8, the number of Quest for Orthologs mammalian proteomes, to 13, the number of mammalian proteomes in the Panther 17 and 18 releases with UniProt release 2023 02. The asterisks (*, **) indicate changes in the dog proteome mapping in release 2023 04 and the horse proteome in 2023 05. For release 2024 01, an additional 22 proteomes were added to the analysis pipeline. (A) proposed changes by proteome; (B) the total number of proposed changes.

**Figure 7:**
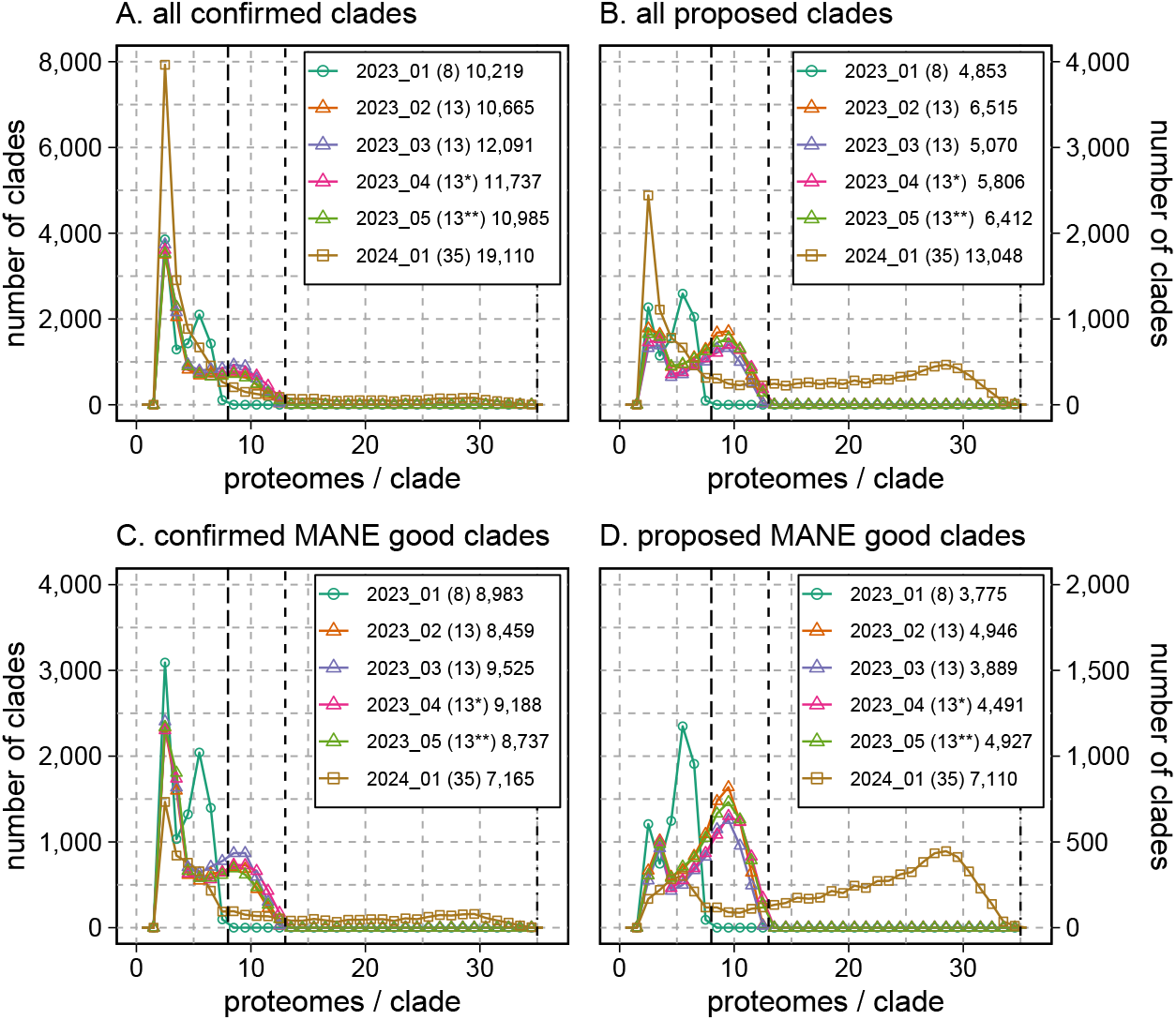
Sizes of clades used to confirm current canonical isoform assignments (A,C), or to propose changes in the canonical isoform (B,D). Each line (color) indicates the UniProt release also plotted in Fig. 6. Numbers in parentheses in the legend number show the numbers of proteomes used to construct the gap-distance trees from which the clades were identified; this is the largest possible size of a clade (also shown with the black dashed vertical lines). The numbers after the maximum clade size indicate the number of orthogroups that had a confirmed (A, C) or proposed (B, D) clade. Asterisks (*, **) indicate changes in the dog proteome mapping and the horse proteome. An additional 22 proteomes were added for release 2024 01. (A, B) all confirmed or proposed clades. (C, D) confirmed or proposed clades that include a human protein that was selected by MANE [13].

Source code for the ortho2tree analysis pipeline, and documentation for its installation, is available from GitHub: github.com/g-insana/ortho2tree.

## Results

### 10–20% of alignments between closely related mammalian proteins have long internal gaps

To illustrate the problem of large alignment gaps in closely related mammalian proteins, canonical mammalian proteomes were compared, with the percent identity and longest gap in the highest scoring alignment displayed in Fig. 1. The mammals shown in Fig. 1 diverged over the past 100 million years (My); for comparison we also aligned two bacterial proteomes at the same evolutionary distance, *E. coli* and *S. typhimurium*, and a pair of yeast proteomes (*S. cerevisiae* and *S. arboricola*) that diverged only 14 My ago, but have very similar distributions of protein sequence identity (Fig. 1). The distribution of bacterial alignment identities (ecoli:salty, 120 My) and yeast (yeast:sacar, 14 My) tracks to the human:bovin (94 My), as well as the human:mouse and human:rat (87 My) identities. As expected, comparisons for more closely related mammals (human:gorgo, *G. gorilla*, and mouse:rat) shift the identity distributions higher; almost 77% of the mouse:rat alignments are more than 90% identical, as are 97% of the human:gorgo alignments.

While the mammalian percent identity distributions at *∼*90 My look very similar to the bacterial and yeast distributions, the gap length distributions are very different (Fig. 1B,C). In Fig. 1B, the longest gap between the bacterial sequences spans 8 amino-acids; for the yeast pair the longest gap is 26 residues. In contrast, among the mammalian *>*90% identical alignments, gap-lengths spanning 1,000– 2,000 residues are found. Nonetheless, 60–80% of the mammalian *>*90% alignments have only one or two gaps. Thus, there seem to be two classes of mammalian alignments: *∼*80% with very short gaps that look like the bacterial and yeast alignments, and *∼*20% with much longer gaps. We believe this second class of sequences, with long alignment gaps, reflects incorrect choices of canonical isoforms.

As proteins diverge from a common ancestor, sequences change by accumulating mutations, including both amino-acid substitutions and insertions/deletions. When both bacterial and mammalian proteins have diverged to be 50–60% identical, about half of the alignments have gaps *≥*5 residues (Fig. 1C). And, as expected, in bacterial alignments the number of *≥*5 residue gaps drops sharply as alignment percent identity increases, so that fewer than 1% of *>*90% identical alignments have five or more gaps. A similar pattern is seen with the yeast protein alignments, with *>*90% identity alignments having *≥*5 gaps about 5% of the time. While mammalian alignments have about the same fraction of gaps as yeast proteins from 50 – 80% identity; at higher identities, mammalian alignments have 3–4 times as many *≥*5 gaps, and about 7–10 times as many *≥*10 gaps (bacterial alignments having no *≥*10-residue gaps, 1.3% of yeast *>*90% identical alignments have *≥*10residue gaps, and 7–13% of mammalian alignments have *≥*10-residue gaps). The data shown in Fig. 1C includes all of the *>*90% identity alignments, including the 50–80% of mammalian alignments without gaps. We suggest that the difference between the mammalian gap fractions and the yeast gap fractions in Fig. 1C reflects incorrect canonical isoform choice, rather than a fundamental difference between bacterial or yeast and mammalian protein structural constraints.

#### Isoform selection by ortho2tree

As outlined in the Materials and Methods, the ortho2tree pipeline (Fig. 2) identifies clades of orthologous proteins that align from end to end, with very few gaps. Although we stress the problem of excess internal gaps in Fig. 1 and below in Figs. 4–5, the ortho2tree pipeline assigns costs to both internal and end-gaps, so that members of the clades it selects have very similar lengths. Fig. 3 shows three examples of clades identified by ortho2tree.

Fig. 3A shows a straight-forward example. In UniProt release 2022 05, the cow (BOVIN, A0A3S5ZP43, 210 aa) and opossum (MONDO, A0A5F8GE93, 256 aa) canonical isoforms of Beta-crystallin A4 were considerably longer than the canonical proteins in the other six Quest for Ortholog mammals (196 aa). In this case, both the cow (BOVIN) and opossum (MONDO) suggested canonicals have N-terminal extensions consistent with the 196 aa clade; the basis for these extensions can be seen in the multiple sequence alignment to the right of the phenogram. Fig. 3A shows a clade with zero cost; all the members of that clade align without gaps. (The sequences in that clade will have many differences, but the gap-cost strategy counts only differences caused by gaps.) In the analysis of UniProt release 2022 05, about half of the clades for both the confirmed canonicals and those with proposed changes have zero cost. The remaining half of the clade costs are relatively uniformly distributed between zero and 0.02, the threshold used to accept a low-cost clade (Suppl. Fig. 1).

Fig. 3B presents a more complex canonical-isoform selection. Here, the selection proposed by ortho2tree disagrees with the MANE canonical assignment. In Fig. 3B, ortho2tree finds a low (but not zero) cost clade containing isoforms from all six of the organisms represented in the Panther orthogroup, but where only two of the proteins in the clade (sp|Q80X89|MOUSE and sp|P36510|RAT) were selected as canonicals in UniProt release 2022 05. However, all six organisms have an isoform with a length of 527–530 aa, and the clade containing those isoforms is selected. This clade does not contain the 527 aa human isoform P0DTE4, which is selected by MANE; while that isoform is the same length as the proposed P0DTE4-1 isoform, it uses a very different set of exons, which are shared only in great apes. This example shows the advantage of multiple sequence alignment and tree-construction; simple length comparison would not be able to discriminate between the two isoforms.

The ortho2tree clade-finding algorithm seeks to identify all of the low-cost clades with nonoverlapping sets of organisms and genes. Thus, in Fig. 3A, the tr|A0A3S5ZP43|BOVIN is highlighted as the incorrect canonical, rather than sp|P11842|BOVIN, because the former protein belongs to a Gene-Centric group that contains tr iso|Q6DTZ8|BOVIN, which can be placed in the correct clade. (Both tr|A0A3S5ZP43|BOVIN and sp|P11842|BOVIN are included in the multiple sequence alignment because they both belong to the PTHR11818:SF19 orthogroup.) In contrast, sp|P11842|BOVIN is the only member of its GeneCentric group. Suppl. Fig. 2 shows a case where three low-cost clades are found in 38 sequences from 7 organisms. Two of the clades, with sequences from 6 (P1) and 7 (P2) organisms, propose canonical isoform changes; the third confirms canonical sequences from 5 of the 7 organisms; and each of the three clades contains a MANE-select human isoform. By building a multiple sequence alignment, a gap-based evolutionary tree, and recognizing different Gene-Centric groups, ortho2tree can make better informed canonical selections.

### Ortho2tree confirmed-canonical isoform alignments are similar to bacterial and yeast alignments

While Fig. 1B indicates that 20–30% of mammalian pairwise alignments have maximum gap lengths that are considerably higher than those seen in bacterial alignments, 80–90% of the alignments have gaps shorter than five residues, and 60–80% of alignments have no gaps. This is consistent with the observation (Table 1) that about 70% (11,010 / 16,015) of the Panther orthogroups produce low-cost clades that contain only sequences currently selected as canonical (confirmed-canonical). The sequences in the “confirmed” ortholog groups tend to be slightly more similar to each other than the complete proteomes (Fig. 4); 100% of the “confirmed” human:gorgo alignments are more than 90% identical, up from 97% in Fig. 1A, and 56% of the human:rodent alignments are more than 90% identical, up from 43% for the complete proteome. (The distribution of bacterial and yeast alignment identities in Figs. 1, 4, and 5 are identical, because exactly the same alignment data is being plotted in the three figures.)

When the “confirmed” mammalian alignments are summarized with respect to percent identity, maximum gap-length, and number of gaps *≥*5, their gap frequencies are similar to those found in bacterial protein alignments and yeast protein alignments (Fig. 5). The bacterial and yeast alignments are useful reference points, because the bacterial proteomes do not have isoforms, while the yeast proteomes have very few (31 out of 6,060 for *S. cerevisiae*, 6 out of 3,653 for *S. arboricola*). For these proteomes, gaps in these alignments must reflect natural structural variation.

Unlike the complete proteome alignments in Fig. 1B, where more than 1,100 alignments have gaps *≥*50 residues, the longest internal gap in the “confirmed” canonical set is 39, and there are only 74 (human:bovin) – 119 (human:gorgo) *>*90% identical alignments with gaps *≥*5 (Fig. 4C). These gap frequencies are 10–20-times lower that those seen in the complete proteome alignments (Fig. 1C). That the “confirmed” orthologous mammalian proteins have a distribution of gap statistics that lies between the bacterial and yeast distributions both supports the argument that correctly annotated canonicals have gap distributions that look like alignments of bacterial and yeast proteins that do not have isoforms, and suggests that the ortho2tree pipeline reliably selects canonical sequences.

### Ortho2tree suggestions reduce alignment gaps in closely related proteins

In complete mammalian proteomes, 13–19% percent of alignments between closely related proteins (*>*90% identical) have gaps longer than 5 residues, 8–13% have gaps longer than 10 residues, and the longest gaps can be more than 1000 residues (Fig. 1B, red rectangle). This contrasts sharply with bacterial and yeast proteins with similar protein identity profiles. As noted earlier, less than 1% of *>* 90% identical *E. coli* vs *S. typhimurium* alignments have gaps *≥*5 residues; for *S. cerevisiae* vs *S. arboricola*, about 5% of *>*90% identical alignments have *≥*5-residue gaps. The ortho2tree pipeline proposes alternative isoforms that allow closely related proteins to align without long gaps.

The effects of the isoform selection on gap length and gap number is summarized in Fig. 5, which shows that while the changed “canonical” and “proposed” isoforms have similar distributions of percent identity (Fig. 5A vs D), the maximum gap length, and the fraction of sequences with gaps *≥*5 drops dramatically in alignments of the “proposed” isoforms (Fig. 5E,F). Like the “confirmed” canonical isoforms (Fig. 4), the “proposed” isoform alignments have gap lengths and gap frequencies that lie between the bacterial and yeast distributions (Fig. 5E,F). By identifying low-gap-cost clades, ortho2tree selects sets of canonical proteins with gap distributions that are very similar to gap distributions from proteomes that do not contain isoforms.

#### ortho2tree **suggests shorter isoforms**

Prior to the implementation of the ortho2tree pipeline, for unreviewed (UniProtKB/TrEMBL) protein sequences, the longest sequence in the GeneCentric group was chosen as canonical. Starting in 2023, UniProt is using the ortho2tree pipeline to select UniProtKB/TrEMBL canonicals on an increasing number of proteomes. As illustrated by the examples in the trees in Fig. 3, and Suppl. Figs. 2 and 3, this rule means that most proposed canonical sequences will be shorter than the previous canonical assignment of unreviewed UniProtKB/TrEMBL entries. Suppl. Fig. 3 summarizes the length changes from canonical to proposed canonical for the changes to the five proteomes highlighted in Fig. 1 in the analysis of UniProt release 2022 05 data. As expected, the changes involving UniProtKB/TrEMBL (unreviewed) proteins (Suppl. Fig. 3B) produced shorter canonical proteins, with a mode change in length of about 30 residues.

When changes involving UniProtKB/Swiss-Prot (reviewed) proteins are included (Suppl. Fig. 3A), the ortho2tree pipeline suggests that hundreds of canonical human and mouse proteins may be too short. In these cases, the shorter canonical isoform in UniprotKB/Swiss-Prot may have been selected by curators because of a publication that references that specific isoform, but ortho2tree selects a longer isoform because it is more similar to other orthologues. The ortho2tree pipeline does not change UniProtKB/Swiss-Prot (reviewed) canonical choices; those suggestions are evaluated individually by curators.

### Ortho2tree adjusts 5–10% of mammalian canonical protein selections

The ortho2tree pipeline was first used to modify canonical isoform selection from the eight Quest for Ortholog mammals (human, chimpanzee, gorilla, mouse, rat, cow, dog, opossum) in the UniProt 2023 01 release of 22 February 2023. Fig. 6 shows all the changes proposed from February 2023 – January 2024; only changes involving unreviewed (UniProtKB/Trembl) sequences were automatically applied to the released reference proteome. With release 2023 02, ortho2tree was applied to the five additional mammals annotated by the Panther ortholog database.[22] The increase in canlf (dog) suggestions reflects a change in canlf orthogroup mapping; Panther 17 (used for releases 2023 01 to 2023 05) used an older version of UniProt with a different set of dog protein accessions. For 2023 03, we constructed a custom ortholog mapping by transferring human Panther orthogroup accessions to dog proteins that were at least 50% identical and aligned over more than 75% of the human sequence, which increased the number of dog suggestions from 19 to 1291. The increase in horse suggestions reflects the change from Ensembl 108 to 109, which produced a different set of horse proteins. Increases from 2023 04 to 2023 05 are due to a shift to Panther 18, which included a new set of dog proteins. Increases in 2024 01 are due to the use of a larger proteome set that included 22 additional mammals, which were assigned to orthogroups by mapping to human proteins using the 50% identity/ 75% coverage rule, as was done earlier with dog.

Once a new proteome is included in the analysis, the number of proposed changes remains relatively constant, except when there are intervening changes in Panther mappings or a new Ensembl genome version. During 2023, the number of suggestions per proteome varied from 500–3,000. (The larger numbers in Fig. 6 reflect the cumulative nature of the plot, changes in ortholog mappings and Ensembl versions produced multiple changes to the same dog and horse proteins.) The number of suggested changes does not correlate with BUSCO [28] proteome quality scores; non-human primates (gorilla, chimpanzee, and macaque) have the largest number of proposed changes. However, across the set of mammals we have examined, 2,000 changes are proposed on average, about 10% of the 20,000 proteins per mammalian proteome.

#### ortho2tree clade sizes

The ortho2tree pipeline does not require that every proteome be included in a low-cost clade; for the releases between 2023 01 and 2024 01, it required only three organisms to be present. Even with that modest requirement, at each release 3,000–5,000 orthogroups (25%) are not considered because they contain sequences from fewer than three proteomes. Fig. 7 shows how the sizes of the “confirmed” and “proposed change” clades have increased over time. For all the releases, there is a peak at three proteomes, and a second at 70–80% of the maximum clade size.

Ortho2tree ranks clades using several criteria: the clade cost, the number of canonical sequences in the clade (with higher weightings for canonicals from human, mouse, and rat), the number of UniprotKB/Swiss-Prot sequences in the clade, and the difference in the lengths of the proposed versus canonical sequences (clades with small changes in length are ranked lower). Thus, proposed clades often do not have sequences from every proteome examined. The weighting system allows ortho2tree to focus on alternate isoform selections that dramatically reduce the length discrepancies (and thus the number of alignment gaps) between the orthologous isoforms. Perhaps because of the evidence ranking required by ortho2tree to select low-cost clades, “proposed change” clades tend to have more organisms than “confirmed” clades.

### Comparison to MANE and APPRIS

For clades that involve human proteins, ortho2tree canonical suggestions generally agree with the selections of the MANE consortium [13]. For the analysis of UniProt release 2022 05, described here in detail, confirmed clades with human sequences agreed with MANE 92% of the time; for proposed changes, there was 82% agreement. As Fig. 7 shows, focusing on clades where the human selection agrees with MANE (“MANE good” clades) shifts the number of proteomes per clade higher; “MANE bad” clades tend to be smaller, or have less evolutionary support.

We also mapped the human UniProt release 2022 05 confirmed clades and proposed changes to APPRIS [14]. 95% of the human accessions in confirmed clades that were labeled either PRINCIPAL or ALTERNATE were PRINCIPAL; for clades with proposed changes, 81% were labeled PRINCIPAL. 88% of “MANE good” proposed changes were also labeled PRINCIPAL by APPRIS, while about 46% of “MANE bad” proposed changes are APPRIS PRINCIPAL. Thus, there is very high agreement between our confirmed and proposed changes with MANE and APPRIS for human proteins, which suggests that ortho2tree suggestions for other organisms are also reasonable.

## Discussion

Alignments between closely related (*>*90% identical) canonical proteins from higher organisms contain a surprising number of long gaps. Here, we argue that these gaps are artifacts of canonical isoform selection. The ortho2tree pipeline identifies clades of orthologous isoforms that align with very few gaps and suggests more biologically reasonable canonical isoforms. Unlike bacterial or yeast protein alignments, high (*>*90%) identity mammalian protein alignments have large numbers of gaps, some of which are hundreds of residues long (Fig. 1). The ortho2tree pipeline suggests alternate canonical isoforms in about one-third of the ortholog families it examines, while two-thirds are “confirmed” to be consistent within the orthologous families (Table 1). The gap-distributions of the “confirmed” proteins are very similar to those of bacterial proteins (Fig. 4) and the changes suggested by the ortho2tree pipeline produce protein sequence alignments with gap distributions that lie between bacterial and yeast alignments (Fig. 5). We consider the bacterial and yeast alignments a reference standard for the kinds of gaps that are expected with sequence divergence—bacterial proteins do not have isoforms, and yeast proteins have very few. Thus, we are encouraged that the isoforms confirmed or proposed by ortho2tree have similar gap properties.

To explore the strengths and weaknesses of the ortho2tree strategy, it is helpful to imagine what the classification results would look like in the ideal case where every mammal has the same set of orthologous genes. Table 1 summarizes the number of Panther orthogroups and canonical proteins that were successfully classified for the first use of ortho2tree in production. The analysis of UniProt release 2022 05 began with 141,544 canonical proteins in 23,878 orthogroups from 8 proteomes. If each of the 8 proteomes contained a canonical protein for each of the Panther orthogroups, we would expect 23,878 *×* 8 = *∼*191,000 canonical proteins; the lower 141,544 number in part reflects the Panther orthogroup subfamily classification; some orthogroup sub-families are found in some of the eight mammals but not in others. Because not every mammal has every orthologous gene, there are some orthogroups that are not found in even 3 mammals (Table 1, “Skipped”) and others where an alignment of the orthologs does not produce a low-cost clade (“No low-cost clades”). And, as Table 1 and Fig. 7 show, even when there are low-cost ortholog clades, they often do not contain a protein from every included proteome. For both confirmed canonical selections and proposed changes, about three-quarters of the canonicals in the orthogroup are found in a low-cost clade (Table 1, “total canonicals” vs “clade canonicals” for both “Confirmed” and “Proposed changes”). For the proteins that are assigned to low-cost clades, about 10% (7,804 / 79,588) are ortho2tree proposed changes. Because automatic changes are made only to UniProtKB/TrEMBL entries, about 85% of the proposed changes were incorporated into the next UniProt release, UP2023 01.

In addition to protein sequences (canonical and isoforms), the ortho2tree strategy requires an ortholog mapping between organisms, and a robust method for finding and ranking low-cost clades. In 2023, ortho2tree analysis began with the 8 Quest for Ortholog mammals (release 2023 01), was extended to the 13 mammals annotated in the Panther ortholog database (releases 2023 02 – 2023 05), and for 2024 01 it was applied to a further set of 35 mammals using a simple ortholog mapping strategy: human Panther assignments were extended to other mammalian sequences that were *≥*50% identical and aligned over 75% of the human protein length. This strategy will allow ortho2tree to be applied to the 132 mammalian proteomes currently in the UniProt Reference Proteome set.

For ortho2tree to make useful canonical suggestions, it must discriminate between alignment gaps that are artifacts of isoform selection, and alignment gaps that result from sequence divergence. Figs. 4 and 5 show the range of sequence identity that ortho2tree has addressed for mammals with our current clade-finding parameters. While the most distant mammalian alignments share as little as 50% identity, alignments from the confirmed clades in Fig. 4 and the proposed changes in Fig. 5 have higher identities, with the most distant alignments dropping to 60% identity. Likewise, the overall distribution of confirmed and proposed clade alignments is more identical; for the complete proteomes, 43% of the human:rodent alignments are *>*90% identical (Fig. 1), while for the confirmed (Fig. 4) and proposed (Fig. 5) orthologs, *∼*58% meet the 90% threshold. This shift to higher identities in part reflects the conservative nature of the ortho2tree pipeline. At higher identities, gaps are produced by isoform differences, while at lower identities, gaps are caused by both isoform differences and evolutionary divergence, so our conservative cost thresholds enrich for higher-identity alignments with isoform-based gaps.

We believe that ortho2tree can also improve canonical selection in more divergent sequence sets, such as vertebrates and plants. Suppl. Fig. 4 shows that human vs *X. tropicalis* (xentr, 340 My) alignments are considerably more divergent than human vs rodent, with about the same divergence as higher plants. At these higher distances, about 50% of alignments are *∼*70% identical, rather than 90% identical. But even at 70% identity, there is a clear separation between the distributions of gaps lengths and frequencies in the “no-isoform” bacterial and yeast alignments, compared with the much longer and more frequent gap distributions in plants and *X. tropicalis*. The gap-distance strategy allows ortho2tree to focus on differences between sequences caused by gaps longer than a threshold, so that short gaps and amino-acid substitutions do not increase the measured distance between two sequences, while highly identical sequences with large gaps (produced by different isoforms) end up in different clades. Traditional sequence difference calculation obscures the difference between distance that reflects evolutionary change and distance that reflects alternative isoform alignment. Longer gap lengths distinguish eukaryotic spliced isoforms from bacterial and yeast proteins, and ortho2tree can exploit those differences to select canonical isoforms more accurately.

The central role of canonical sequences in reference protein databases cannot be overemphasized. Canonical sequences comprise the libraries used for similarity searching, and they form the foundation for virtually all derivative protein analyses and databases. Canonical sequences are used for structure and function prediction, domain annotation, evolutionary rate analysis, and orthology assignment. Our analysis suggests that 10–20% of canonical sequences in well-annotated genomes are likely misassigned; the ortho2tree analysis pipeline reduces those errors so that confirmed and corrected sequences have gaps in the ranges seen in yeast and bacteria. Since the first release of corrected canonical assignments in February 2023, ortho2tree has corrected about 50,000 proteins from 35 mammals, a number that will more than double in May 2024. As ortho2tree is applied more broadly, our ability to characterize proteins structurally and functionally will continue to improve.

## Author contributions statement

W.R.P. conceived the experiments and developed the initial version of the ortho2tree pipeline, G.I. restructured the pipeline for integration into the UniProt Reference Proteome production process and performed extensive testing. G.I., M.J.M. and W.R.P. analyzed the results. G.I., M.J.M. and W.R.P. wrote and reviewed the manuscript.

## Acknowledgments

This work is supported in part by a grant from the National Science Foundation (NSF: #1759626) to W.R.P. and by European Molecular Biology Laboratory core funds.

## Data availablility

The data for the analysis of UP2022 05 QfO proteomes, discussed in the manuscript, and the data and R scripts used for the creation of the figures is available from GitHub (github.com/g-insana/ortho2tree) and from the Zenodo resources: doi. org/10.5281/zenodo.10778115 and doi.org/10. 5281/zenodo.11113232.

## SUPPLEMENTAL FIGURES

**Suppl. Fig. Fig. 1:**
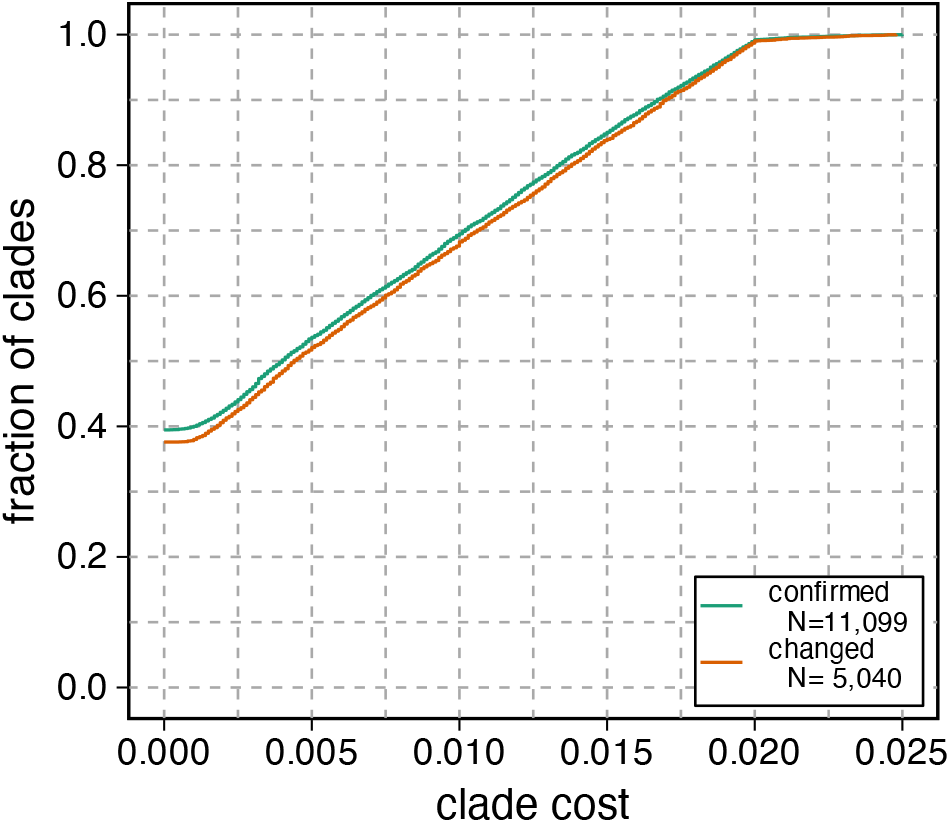
Distribution costs in clades with confirmed or proposed changes in the canonical isoform assignment. The fraction of all confirmed or proposed clades is plotted against the gap-based clade cost. Clades with zero cost contain sequences that align without gaps from beginning to end.

**Suppl. Fig. Fig. 2:**
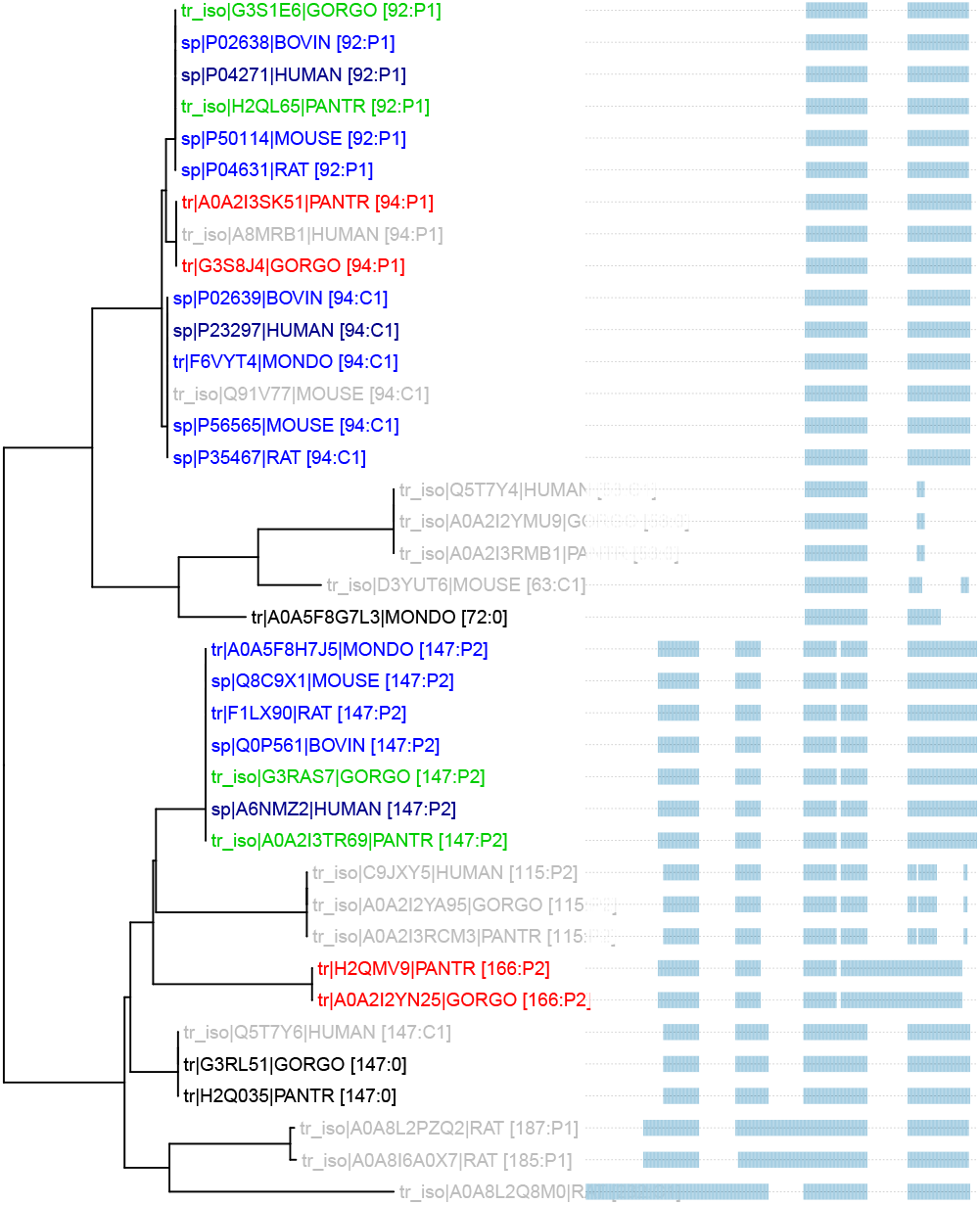
A phenogram showing three clades found for members of the PTHR11639:SF134 ortholog group, that contains S100-A1 small calcium binding proteins. Sequences are labeled as in Fig. 3, but here there are two clades with proposed changes (P1 and P2), as well as a confirmed clade (C1).

**Suppl. Fig. Fig. 3:**
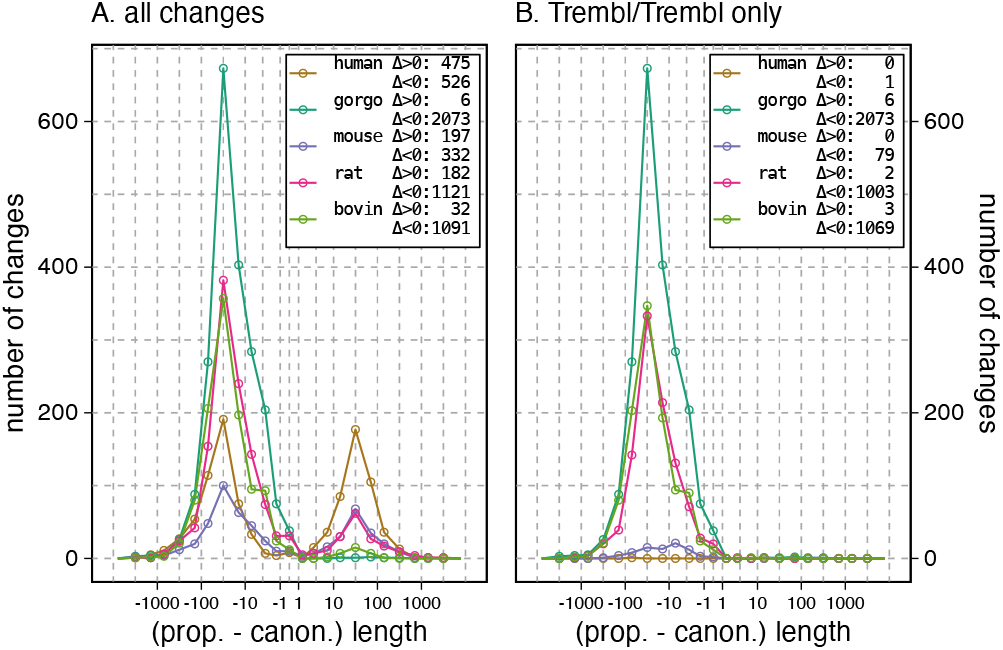
Changes in canonical sequence length for proposed changes. Differences between the proposed and current canonical sequence lengths for five different proteomes are shown. (A) All proposed canonical changes, including SwissProt to SwissProt, SwissProt to TrEMBL, and TrEMBL to TrEMBL; (B) Only TrEMBL to TrEMBL changes. Numbers in the legend box indicate the total number of changes proposed for that organism. Only 11 of the 4,238 proposed Trembl/Trembl changes increase the length of the canonical sequence for these 5 proteomes.

**Suppl. Fig. Fig. 4:**
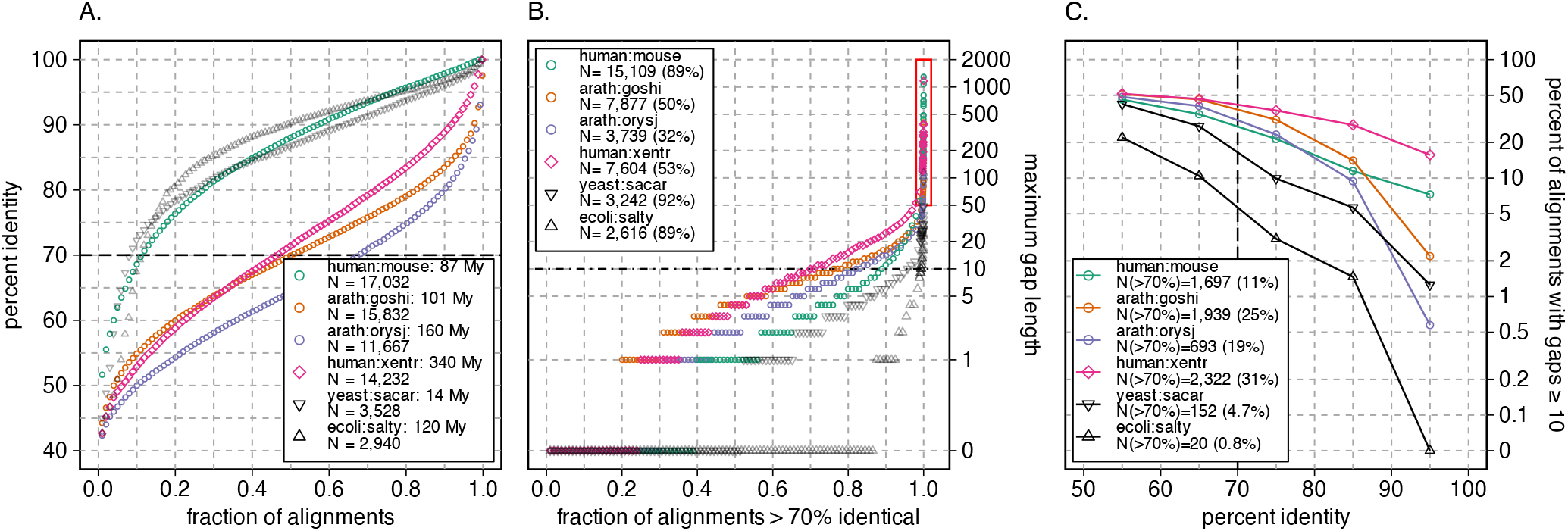
Comparison of sequence identity and gap lengths in more divergent proteins. This figure replots the data from Fig. 1 for human:mouse, yeast:sacar, and ecoli:salty, and adds identity and gap distribution data for three more distant pairs of organisms, human vs *X. tropicalis* (human:xentr, 340 My), and two plant pairs: *A. thaliana* vs cotton (arath:goshi, 101 My) and rice (arath:orysj, 160 My). Panel (A) shows the identity distributions; (B) the maximum gap lengths in alignments that are more than 70% identical, and (C) the number (and fraction) of sequences *>*70% identical with gaps *≥*10.

